# Is maternal thyroid hormone deposition subject to a trade-off between self and egg because of iodine? An experimental study in rock pigeon

**DOI:** 10.1101/2021.01.03.425121

**Authors:** Tom Sarraude, Bin-Yan Hsu, Suvi Ruuskanen, Ton Groothuis

## Abstract

Maternal hormones constitute a key signalling pathway for mothers to shape offspring phenotype and fitness. Thyroid hormones (THs; triiodothyronine, T_3_ and thyroxine, T_4_) are metabolic hormones known to play crucial roles in embryonic development and survival in all vertebrates. During early developmental stages, embryos exclusively rely on the exposure to maternal THs, and maternal hypothyroidism can cause severe embryonic maldevelopment. The TH molecule includes iodine, an element that cannot be synthesised by the organism. Therefore, TH production may become costly when environmental iodine availability is low. This may yield a trade-off for breeding females between allocating the hormones to self or to their eggs, potentially to the extent that it even influences the number of laid eggs. In this study, we investigated whether low dietary iodine may limit TH production and transfer to the eggs in a captive population of Rock pigeons *(Columba livia).* We provided breeding females with an iodine-restricted (I- diet) or iodine-supplemented diet (I+ diet) and measured the resulting circulating and yolk iodine and TH concentrations and the number of eggs laid. Our iodine-restricted diet successfully decreased both circulating and yolk iodine concentrations compared to the supplemented diet, but not circulating or yolk THs. This indicates that mothers may not be able to independently regulate hormone exposure for self and their embryos. However, egg production was clearly reduced in the I- group, with fewer females laying eggs. This result shows that restricted availability of iodine does induce a cost in terms of egg production. Whether females reduced egg production to preserve THs for themselves or to prevent embryos from exposure to low iodine and/or THs is as yet unclear.

## Introduction

Non-genetic inheritance is defined as the transmission of information between generations beyond coding genes (Danchin et al., 2011). Parental effects are included in this non-genetic inheritance, may be considered adaptive (Moore et al., 2019; Mousseau and Fox, 1998; Yin et al., 2019), and parental effect of maternal origin, i.e. maternal effects, have received increasing attention since the 1990’s (Bernardo, 1996; Mousseau and Fox, 1998). Hormones of maternal origin can be transferred to the offspring and constitute a potential pathway for mothers to influence their offspring’s phenotype (Groothuis et al., 2019). Hormone allocation to offspring could be costly for mothers as it could induce a trade-off between allocating hormones to their own metabolism or to their offspring’s. Steroid hormones, the most studied hormones in the context of hormone-mediated maternal effects, may not be that costly to produce as they are derived from cholesterol, which is abundant in the organism (Groothuis and von Engelhardt, 2005). On the other hand, thyroid hormones (THs) may be considered costly, as their molecular structure includes iodine, a trace element that cannot be synthesised by organisms and must therefore be found in the environment.

Thyroid hormones are metabolic hormones that are present in two main forms: thyroxine (T_4_) that contains four atoms of iodine, and triiodothyronine (T_3_) that contains three atoms of iodine. Iodine is concentrated into the thyroid gland and incorporated into tyrosines that will be combined to form T_4_ and T_3_ (McNabb and Darras, 2015). The thyroid gland produces mostly T_4_ and lesser amounts of T_3_. In the peripheral tissues (e.g. liver, kidney, muscle), T_3_ is mostly obtained from T_4_ via removal of an iodine atom by deiodinase enzymes (McNabb and Darras, 2015). TH action mainly depends on TH receptors that have a greater affinity for T_3_ than for T_4_ (Zoeller et al., 2007). This is why T_4_ is mostly seen as a precursor of T_3_, the biologically active form. THs play important roles in growth, reproduction, metamorphosis and thermoregulation, and the TH signalling pathway is well conserved throughout the animal kingdom, from invertebrates (Holzer et al., 2017; Taylor and Heyland, 2017) to vertebrates (reviewed by Ruuskanen & Hsu, 2018).

The main sources of iodine for terrestrial animals is via uptake of food, where plants absorb it from the soil, and via drinking water (Anke, 2004). Because iodine availability differs across environments and food sources (Anke, 2004; Boggs et al., 2011), THs may be costly to produce when iodine availability is limited. Low iodine diet in rodents under laboratory conditions generally decrease circulating THs (e.g. Santisteban et al., 1982; Zhimei et al., 2014; but see Bocco et al., 2020). Furthermore, in wild alligators *(Alligator mississippiensis*) overall higher plasma THs were reported in the populations exposed to higher iodine concentrations compared to low iodine habitat (Boggs et al., 2011; Boggs et al., 2013). In breeding animals, this potential cost may induce trade-offs between allocating resources (i.e., iodine and THs) to themselves or to their progeny because embryos rely exclusively on THs of maternal origin during early development (humans, Stepien and Huttner, 2019; fish, Castillo et al., 2015; birds, Darras, 2019). In humans, TH deficiency arising from iodine deficiency can severely impair brain development and increase foetus mortality (reviewed by Zimmermann, 2009). Although the importance of THs has long been acknowledged, such a potential trade-off has never been studied.

Birds provide an excellent model organism to study hormone-mediated maternal effects and the associated trade-offs (Groothuis et al., 2019). Recent studies in birds have shown that physiological variation in prenatal THs can affect hatching success, growth and metabolism (Hsu et al., 2017; Hsu et al., 2019; Ruuskanen et al., 2016; Sarraude et al., 2020a). In addition, avian embryos develop in eggs, outside of mothers’ body, which facilitates the measurement of not only maternal physiological status but also the actual embryonic exposure to maternal hormones.

Three older studies investigated the effect of different dietary iodine concentrations on the thyroid function in ring doves (*Streptopelia risoria*) and in egg laying Japanese quails *(Coturnix japonica)* (McNabb et al., 1985a; McNichols and McNabb, 1987; Spear and Moon, 1985), and on Japanese quail embryos and hatchlings (McNabb et al., 1985b). McNabb and colleagues (1985a) found that circulating and yolk iodine were proportional to dietary iodine. Thyroid T_4_ was decreased by restricted dietary iodine but plasma THs were not affected in quails (McNabb et al., 1985a). In contrast, McNichols and McNabb (1987) found that both circulating iodine and T_4_ concentrations decreased in response to low dietary iodine in the ring doves. In quails, embryos and hatchlings from the eggs with low iodine concentrations suffered from thyroid gland hypertrophy, but their circulating TH levels were not different from those from the eggs with moderate or high yolk iodine concentrations (McNabb et al., 1985b). However, none of these studies measured yolk THs, which is necessary information to investigate the potential trade-offs between mothers’ circulation and embryonic exposure to THs.

The aim of our study was two-fold. First, to investigate whether limited iodine may constrain TH production, and second, whether mothers in that case can prioritise TH allocation to either self or their eggs. To this end we provided breeding pairs of rock pigeons *(Columba livia)* either with a diet restricted in iodine (hereafter I-), or a diet supplemented with iodine (hereafter I+). If restricted dietary iodine limits TH production, we may expect a decrease in circulating THs in the mother, as found in ring doves, and/or a decrease in yolk THs. To cope with limited TH production due to iodine limitations, mothers may have developed regulatory mechanisms allowing them to trade off THs for self or for the egg. This would be reflected in a different distribution of TH concentration between mothers’ circulation and in the egg relative to controls. Alternatively, mothers may prioritise circulating and yolk TH concentrations by limiting egg production and thereby the total amount of TH allocated to yolks, thus prioritising quality over quantity of offspring. This would be consistent with two studies that showed that hypothyroidism ceased egg laying in Japanese quails (Wilson and McNabb, 1997), and reduced egg production in chickens (Van Herck et al., 2013). We also expected the effect of iodine restriction to be more pronounced as the exposure duration to the treatment becomes longer. For example, iodine stores may deplete with time in the I- diet, thus decreasing circulating and/or yolk iodine and THs. In addition, laying several clutches under limited iodine availability may further deplete iodine and TH stores. We expected therefore yolk iodine and THs to decrease with clutch order in the I- diet. In general, and in line with previous studies (McNabb et al., 1985b; McNichols and McNabb, 1987), we may expect limited iodine to have a stronger effect on T_4_ than on T_3_, as T_4_ needs one more atom of iodine and is much less biologically active.

## Material and methods

### Study species and housing conditions

The experiment was conducted in 2018 on 38 pairs of wild-type rock pigeons *(Columba livia).* Rock pigeons lay two eggs per clutch, with a 48-hour interval between the two eggs. In addition, rock pigeons can lay multiple clutches in a single breeding season (Johnston et al., 1995). The birds were identified by unique ring code and combination, and were housed in a large outdoor aviary (45 m long x 9 m wide x 4 m high) in Groningen, the Netherlands, divided in 4 equal compartments (2 compartments per treatment, n = 9–10 breeding pairs per compartment, see below). The aviary included enough nest boxes and nesting material for all the breeding pairs. Before the experiment, all birds were fed a standard diet for pigeons (seed mixture Kasper™ 6721 + seed mixture Kasper™ 6712 + pellets P40 Kasper™ 6700). Standard food, water and grit were provided ad libitum. Before the experiment, 18 eggs were collected from unidentified females under standard diet and used for analyses of yolk iodine (see statistical section below).

### Experimental design

#### Iodine restricted and supplemented diet

We provided the experimental birds with either an iodine restricted (I-, n = 19 pairs) or an iodine supplemented (I+, n = 19 pairs) diet until all eggs and blood samples were collected. Egg collection was ended around three weeks after the initiation of second clutches, leading to a total of approx. 10 weeks (see below for more details). The restricted diet contained 0.06 mg of iodine/kg of food (Altromin™ C1042) and the supplemented diet was the same food supplemented with 3 mg of iodine/kg by the manufacturer. Therefore, both diets had exactly the same composition of all essential micro- and macronutrients except iodine. The restricted treatment corresponds to about 10% of the iodine content in the standard pigeon diet (0.65 mg/kg), and approximately 20% of the minimum iodine requirement for ring doves (0.30 mg I/kg) according to Spear and Moon (1985). In addition, this restricted treatment corresponds to a low iodine treatment (0.05 mg/kg) used in a previous experiment on Japanese quails (McNabb et al., 1985a) that induced a significant decrease in circulating and yolk iodine. The supplemented treatment (3 mg/kg) corresponds to five times the concentration of iodine in the standard pigeon diet. In McNabb and colleagues (1985a), the maximal dietary iodine (1.2 mg I/kg feed) was ca. eight times the sufficient iodine concentration required for Japanese quails (0.15 mg I/ kg feed) and the authors observed no detrimental effects of this high dose. Therefore, we expected no detrimental effect of our supplemented treatment either. Food, water, and grit were provided ad libitum throughout the experiment. We collected both eggs from first and second clutches of females fed with restricted or supplemented iodine diets. We also collected blood samples after clutch completion from the two experimental groups (I- and I+). The second set of samples (i.e. eggs and blood) was collected to test for the effect of exposure duration of the treatment (see timeline below).

#### Timeline of the experiment

Nest boxes were opened and nesting material was provided two weeks after the experimental diet was introduced to stimulate egg laying. Egg laying usually starts within a week of nest-box opening. Based on Newcomer (1978), who fed hatchling chicken with low iodine diet (0.07 mg I/kg feed), we could expect thyroid iodine content to be the lowest from 10 days onwards after introducing the experimental diet. The first eggs (i.e. from the first experimental clutches) were collected 3 weeks after the introduction of the experimental diet and were collected over 12 days. On average, the eggs from 1^st^ clutches were laid 26.4 days (SD = 2.9) days after the onset of the experimental diet. Freshly laid eggs were collected, replaced by dummy eggs to avoid nest desertion. Second clutches were initiated by removing dummy eggs approximately 2 weeks after the completion of the first clutch (i.e., 5 weeks after the start of the experimental diet), and eggs collected over a period of 18 days. On average, the eggs from 2^nd^ clutches were laid 53.5 days (SD = 3.3) after the onset of the experimental diet. We also collected some late first clutches (on average 52.6 (SD = 6.5) days after the onset of the experimental diet).

#### Egg and blood sample collection

Table 1 summarises the number of samples collected. Eggs and blood samples were collected in the exact same manner in the first and second clutches. Freshly laid eggs were collected and were stored in a –20°C freezer. Not all females laid complete clutches of 2 eggs, and several females did not lay an egg at all. Females were captured during incubation in the nest boxes, and blood samples (ca. 400 μl) were taken from the brachial vein after clutch completion (average (SD) = 4 (4.7) days after clutch completion, range = 0–21 days). Unfortunately, we could not blood sample the females that did not lay eggs, as this would have caused serious disturbance to all the birds in the same aviaries as we had to catch them by hand netting in the large aviary. Half of the blood sample (ca. 200 μl) was taken with heparinised capillaries for plasma extraction (for TH analyses) and stored on ice until centrifugation. The other half of the sample was taken with a sterile 1 ml syringe (BD Plastipak ™) and let to coagulate for 30 min at room temperature before centrifugation for serum extraction (for iodine analyses). Previous studies measured iodine in serum samples (McNabb et al., 1985a; McNichols and McNabb, 1987), therefore we decided to measure iodine in the serum for comparable results. Whole blood samples were centrifuged at 3 500 RPM (ca. 1164 G-force) for 5 min to separate the plasma from red blood cells (RBCs), and at 5 000 RPM (ca. 2376 G-force) for 6 min to separate the serum from RBCs. After separation, all samples (plasma, serum and RBCs) were stored in a –80°C freezer for analyses of THs and iodine.

**Table 1:** Summary of the egg and blood samples collected.

|  |  | Untreated | I- |  | I+ |  |
| --- | --- | --- | --- | --- | --- | --- |
|  |  | 1<sup>st</sup><br>clutches | 1<sup>st</sup><br>clutches | 2<sup>nd</sup><br>clutches | 1<sup>st</sup><br>clutches | 2<sup>nd</sup><br>clutches |
| Egg samples | Complete clutches | NA | 6 | 6 | 12 | 5 |
|  | Incomplete clutches | NA | 5 | 2 | 6 | 2 |
|  | Total number eggs | 17 | 17 | 14 | 30 | 12 |
| Blood samples | Number of females | NA | 10 | 8 | 14 | 7 |

6 females could not be captured and sampled (1 in I- and 5 in I+ group).

### Hormone and iodine analyses in plasma and yolk samples

Eggs were thawed, yolks separated, homogenised in MilliQ water (1:1) and a small sample (ca. 50 mg) was used for TH analysis. Yolk and plasma THs were analysed using nano-LC-MS/MS, following Ruuskanen et al. (2018; 2019). TH concentrations, corrected for extraction efficiency, are expressed as pg/mg yolk or pg/ml plasma.

Yolk and serum iodine (ICP-MS, LOD of 3 ng/g of yolk and 1.5 ng/ml of serum) analyses were conducted by Vitas Analytical Services (Oslo, Norway). Yolk iodine was measured in a sample of ca. 1 g of yolk, and serum iodine was measured in a sample of ca. 0.2 ml of serum.

### Statistical analysis

#### General information

Data were analysed with the software R version 4.0.2 (R Core Team, 2020). To test for the effect of iodine restriction on egg laying, we compared the number of females that laid first clutches in both groups, and the total number of eggs laid in first clutches with two Pearson’s chi-squared tests. The rest were fitted with linear mixed models (LMMs) using the R package *lme4* (Bates et al., 2015) and p-values of the predictors were calculated by model comparison using Kenward-Roger approximation with the package *pbkrtest* (Halekoh and Højsgaard, 2014). The response variables were plasma THs concentrations (T_3_, T_4_), serum iodine, and concentrations of yolk THs and yolk iodine. Relevant interactions between predictors were tested by adding them one-by-one in the models with main effects only. Post-hoc tests of interactions were performed with the package *phia* (de Rosario-Martinez, 2015). Model residuals were inspected for normality and homogeneity with the package *DHARMa* with 1,000 simulations (Hartig, 2020). When either of the assumptions was violated, the response was ln-transformed (see Tables) and in these cases the model residuals showed the required distributions. Estimated marginal means (EMMs) were calculated from the models using the package *emmeans* (Lenth, 2019). When the response was transformed, the EMMs were calculated on the back-transformed data.

Although the treatment started for all females on the same date, each female, at the time of egg laying, was exposed to the experimental diet for different durations because of the varying laying dates between females. This may influence the effects of iodine manipulation on circulating and yolk iodine and THs. However, because the second clutches were laid after a longer exposure to the treatments than were first clutches (average exposure duration (SD) 2^nd^ clutches = 53.5 (3.3) days; 1^st^ clutches = 30.9 (10.6) days), clutch order and exposure duration (i.e. the number of days between the onset of the experimental diet and laying date) were confounded. Therefore, we used two different models, one with exposure duration and another one with clutch order. We controlled for egg order (i.e. first or second egg in a clutch) in our models for yolk THs initially since a previous study in the rock pigeons showed a non-significant trend for higher yolk T_3_ concentrations in the second eggs (Hsu et al., 2016). However, we detected no such effect in our models (all *F* < 0.57, all p-values > 0.45) and thus egg order was excluded from the final models.

#### Model specification

Circulating iodine (ln-transformed) and T_3_ and T_4_ concentrations were analysed by fitting an LMM with treatment (I- or I+), exposure duration, completeness of a clutch as a categorical variable (complete or incomplete, i.e., to further test for the effect of number of eggs laid), and the two-way interactions between treatment and exposure duration or completeness as fixed factors. Female identity (for iodine and T_4_, but not T_3_ because of singularity: variance estimate collapsed to 0) and hormone extraction batch (for T_3_ and T_4_) were added as random intercepts.

Yolk components (iodine, T_3_, T_4_) were analysed in two different sets of models. The first set of models, for yolk iodine only, compared untreated eggs (collected before the start of the treatment) to the eggs collected in the two treatments (I+ and I-). This way, we could test whether yolk iodine differed between untreated and experimental eggs. Here we only used experimental first clutches as only one clutch of eggs per untreated female was collected. The model only included treatment (a three-level categorical predictor: untreated, I+ and I-) as the predictor.

The second set of models tested the effect of iodine treatment, exposure duration to the treatment or clutch order on yolk iodine and THs. Here we included both 1^st^ and 2^nd^ clutches from iodine treatments, but no eggs from untreated females. Yolk iodine (ln-transformed) was first analysed in a LMM that included treatment as a categorical variable, exposure duration (days since the start of the experiment), completeness of a clutch (complete or incomplete) and the two-way interactions between treatment and exposure duration or completeness, and female identity as a random intercept. This LMM somewhat violated the assumption of homogeneity of variances between the groups because of the larger variance in yolk iodine in the I+ group. Nevertheless, such a violation should not undermine our results as a recent paper demonstrated that LMMs are fairly robust against violations of distributional assumptions (Schielzeth et al., 2020). Yolk iodine was also analysed in a second model that included treatment, clutch order (1^st^ or 2^nd^ clutch, categorical variable) and their interaction as the predictors. Yolk T_3_, T_4_ (ln-transformed) were analysed using the same models (with exposure duration or clutch order) as for yolk iodine. Hormone extraction batch was added as a random intercept for yolk T_3_ and T_4_.

## Results

### Circulating iodine and TH concentrations

In line with our expectations, there was a clear effect of iodine treatment on circulating iodine concentrations: serum iodine was about four times lower in the I- group than in the I+ group (raw data average (SE), I- = 11.1 (1.2) ng/ml serum, I+ = 44.0 (4.7) ng/ml serum; Table 2, Fig. 1). The effects of clutch completeness, exposure duration and their interactions with treatment on serum iodine were statistically unclear (Table 2).

**Figure 1:**
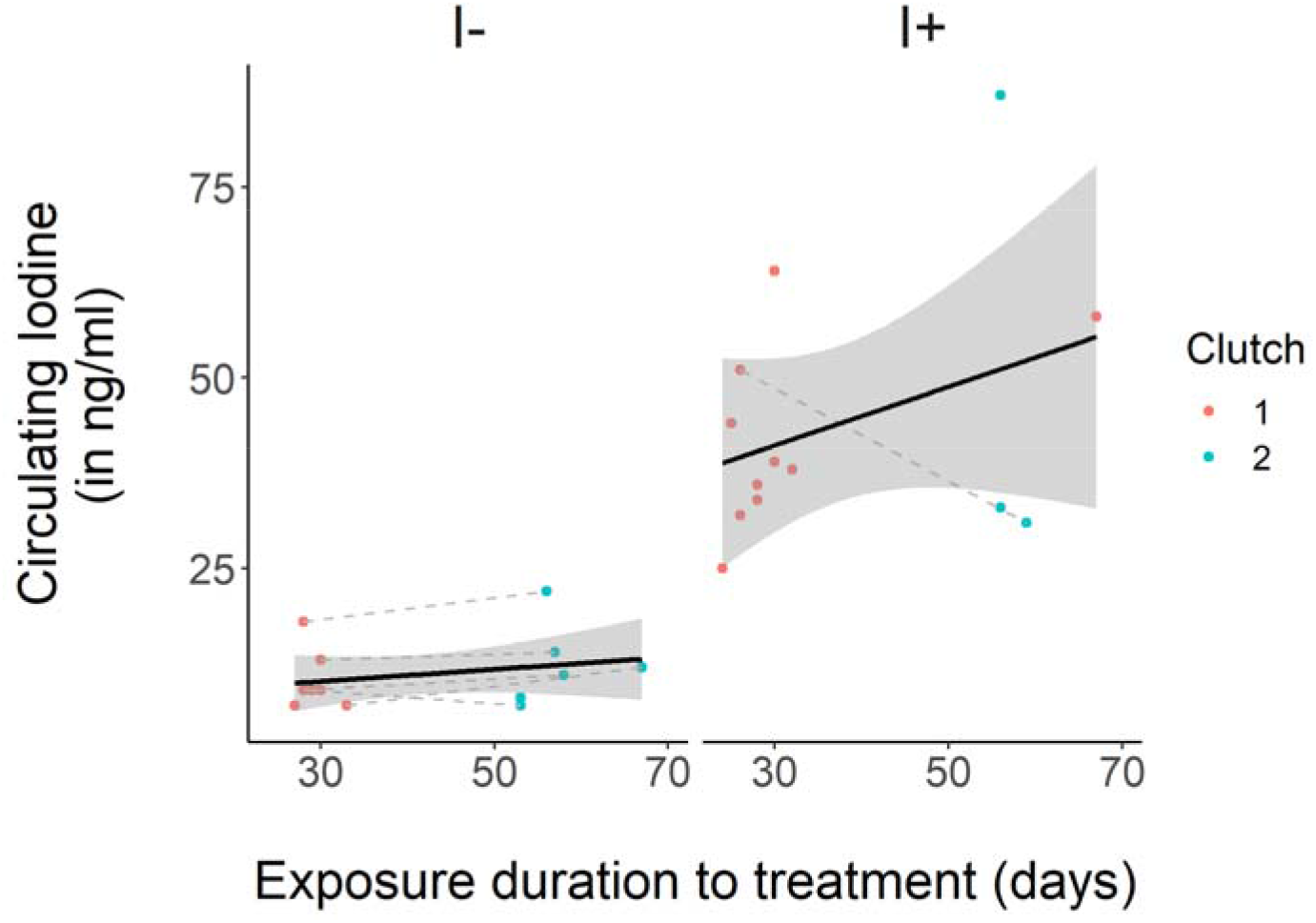
Circulating iodine in rock pigeon females treated with an I- or I+ diet. Black lines and shadow areas represent average and s.e.m values within each group, and grey dashed lines connect blood samples from the same females. Some females were only captured once, hence not all dots are connected. See Table 1 for sample sizes.

**Table 2:** Results of the LMMs on circulating THs and iodine from rock pigeon females treated with an I- or I+ diet.

| Predictor | Ln-Iodine |  |  | T<sub>4</sub> |  |  | T<sub>3</sub> |  |  |
| --- | --- | --- | --- | --- | --- | --- | --- | --- | --- |
|  | Estimate<br>(SE) | F<sub>ddf</sub> | p | Estimate<br>(SE) | F<sub>ddf</sub> | p | Estimate<br>(SE) | F<sub>ddf</sub> | p |
| Treatment (I+) | 1.43<br>(0.15) | 93.6<sub>17.0</sub> | **<0.001** | -0.68<br>(2.53) | 0.07<sub>18.6</sub> | 0.79 | -0.02<br>(0.37) | 0.002<sub>32.0</sub> | 0.96 |
| Exposure duration | 0.005<br>(0.004) | 1.63<sub>12.0</sub> | 0.23 | 0.13<br>(0.05) | 6.65<sub>18.4</sub> | **0.02** | 0.002<br>(0.01) | 0.04<sub>32.0</sub> | 0.85 |
| Complete clutch (Yes) | -0.01<br>(0.15) | 0.01<sub>22.5</sub> | 0.93 | -2.29<br>(2.46) | 0.79<sub>30.1</sub> | 0.38 | 0.35<br>(0.41) | 0.73<sub>32.0</sub> | 0.40 |
| Treatment (I+) * exposure duration | -0.003<br>(0.008) | 0.12<sub>12.0</sub> | 0.73 | 0.19<br>(0.09) | 4.34<sub>15.6</sub> | 0.05 | -0.005<br>(0.02) | 0.04<sub>31.0</sub> | 0.83 |
| Treatment (I+) * Complete clutch (Yes) | -0.06<br>(0.32) | 0.04<sub>21.2</sub> | 0.85 | -2.48<br>(4.93) | 0.23<sub>30.2</sub> | 0.64 | -0.35<br>(0.84) | 0.17<sub>31.1</sub> | 0.68 |

Plasma T_3_ was not affected by supplementation or restriction of iodine, nor by exposure duration (Table 2; Fig. 2A). There was an almost statistically significant interaction between iodine treatment and exposure duration of the treatment on plasma T_4_ (Table 2). A post-hoc test of the interaction showed that plasma T_4_ increased with time in the I+ group but not in the I- group (adjusted slope (SE) I+ = 0.24 (0.07), χ^2^ = 12.9, Holm-adjusted p < 0.001; adjusted slope (SE) I- = 0.05 (0.06), χ^2^ = 0.56, Holm-adjusted p = 0.46; Fig. 2B). Yet, the large confidence intervals warrant due caution in interpreting this trend. There were no clear effects of clutch completeness and its interaction with treatment on plasma THs (Table 2).

**Figure 2:**
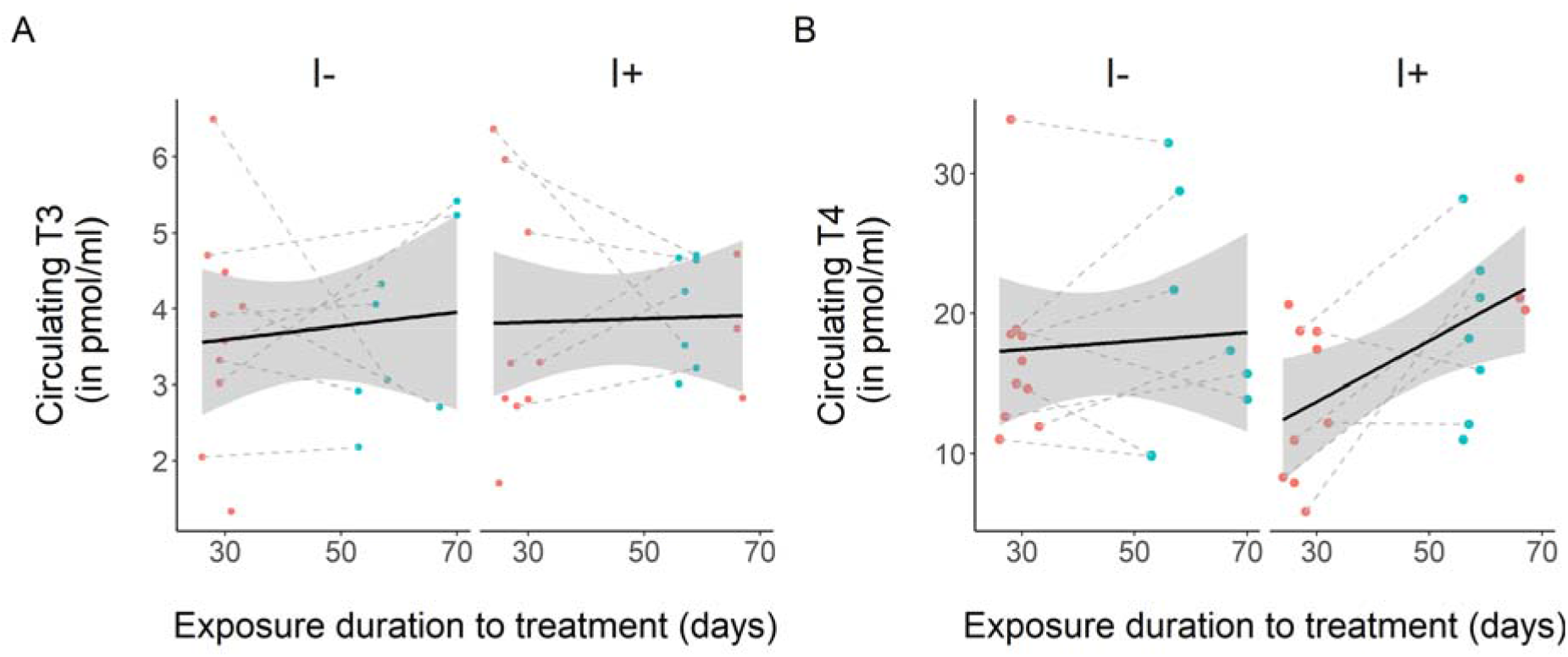
Circulating T_3_ (A) and T_4_ (B) in rock pigeon females treated with an I- or I+ diet. Red dots refer to blood samples after collected the first clutches and blue dots refer to blood samples collected after the second clutches. Black lines and shadow areas represent average and s.e.m values within each group, and grey dashed lines connect blood samples from the same females. Some females were only captured once, hence not all dots are connected. See Table 1 for sample sizes.

### Egg iodine and egg TH concentrations

#### Untreated eggs vs 1^st^ clutches of the iodine treatments

In line with our prediction, eggs from the I- group had ca. seven times lower iodine levels than eggs from the I+ group and two times lower iodine levels than untreated eggs; eggs from the I+ group had ca. four times higher iodine levels than untreated eggs (back-transformed EMMs (SE), untreated = 29.7 (4.1) ng/g yolk, I- = 16.1 (2.0) ng/g yolk, I+ = 123.4 (11.9) ng/g yolk; (overall test: LM, *F* = 91.3, p < 0.001); post-hoc Tukey comparisons, all |t| > 3.38 and all p < 0.004, Fig. 3).

**Figure 3:**
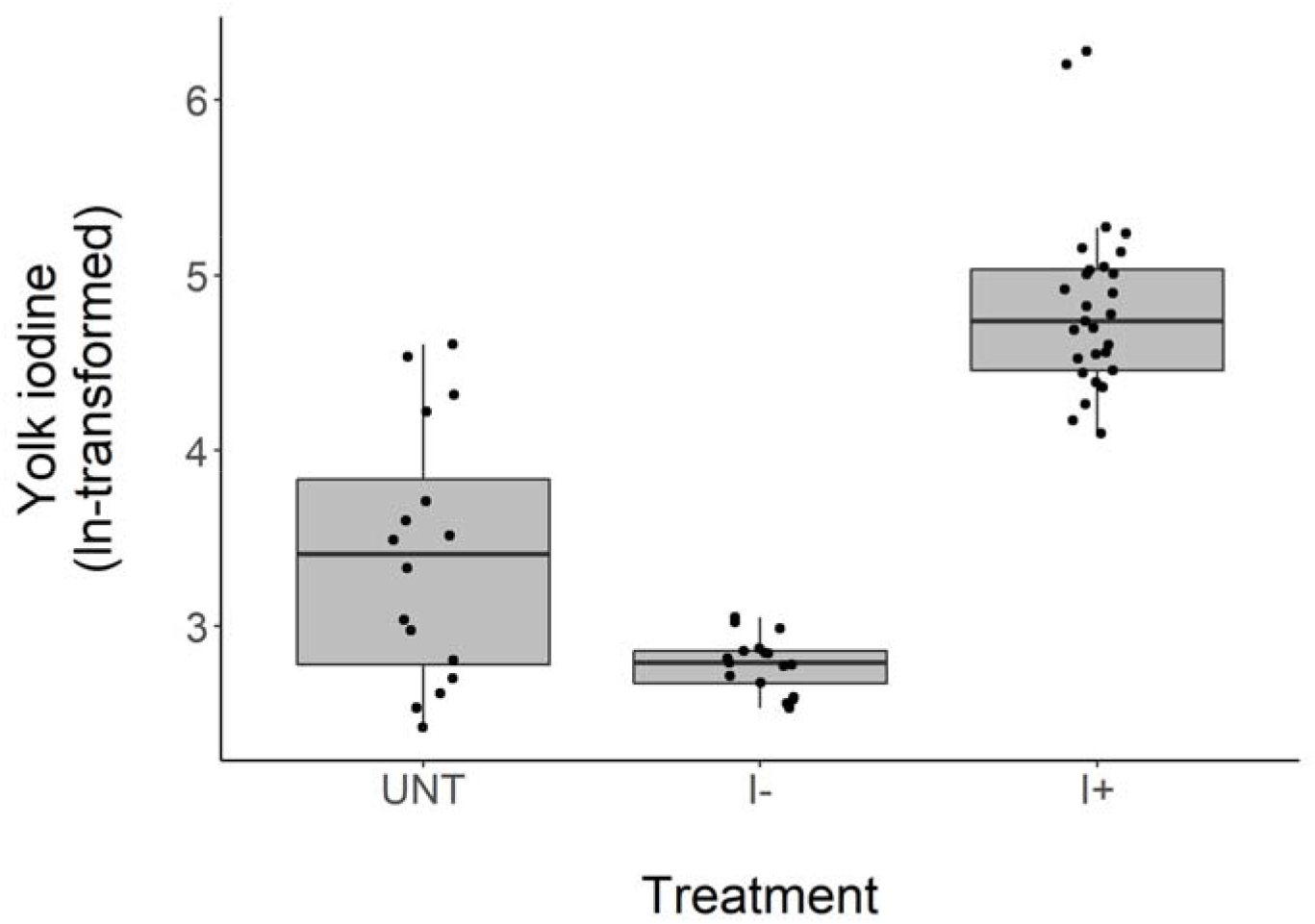
Yolk iodine in eggs from 1^st^ clutches laid by rock pigeon untreated females (UNT), females treated with an I- or an I+ diet. See Table 1 for sample sizes.

#### Effect of exposure duration (experimental eggs from 1^st^ and 2^nd^ clutches)

Also in this dataset, yolk iodine concentration was seven times lower in eggs of I- treated females than in I+ females (mean (SE), I- = 16.03 (0.52) ng/g yolk, I+ = 117.48 (12.36) ng/g yolk; Table 3, Fig. 4), but longer exposure duration or clutch completeness did not influence yolk iodine concentration (Table 3). Yolk iodine tended to be lower in the 2^nd^ clutches, but this difference was not related to dietary iodine (Table 3; Fig. 4).

**Figure 4:**
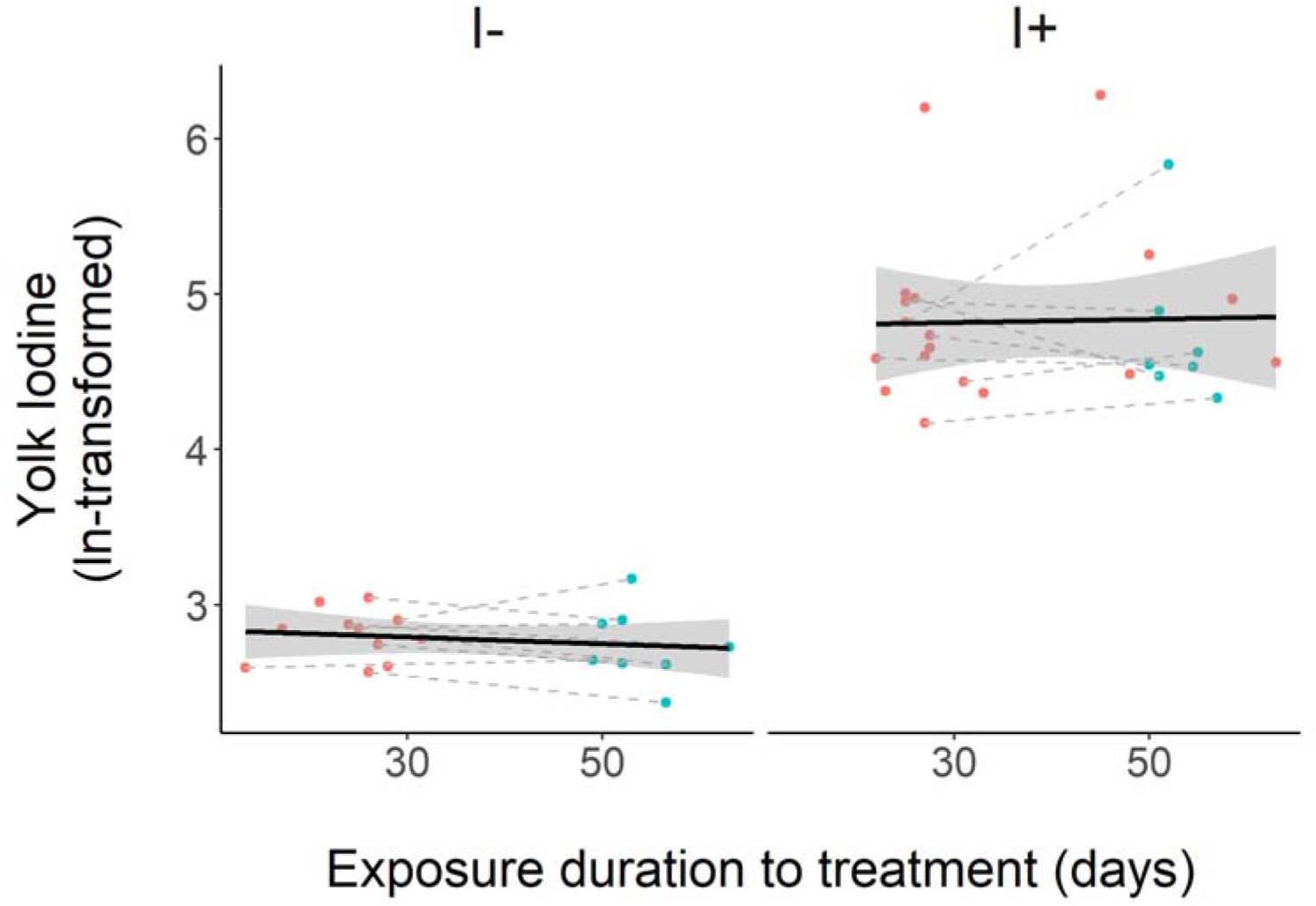
Yolk iodine in eggs from 1^st^ and 2^nd^ clutches laid by rock pigeon females treated with an I- or I+ diet. Eggs from the same female and the same clutch were averaged. Red dots refer to eggs from the first clutches and blue dots refer to eggs from the second clutches. Black lines and shadow areas represent average and s.e.m values within each group, and grey dashed lines connect eggs from the same females. Some females did not lay two clutches, hence not all dots are connected. See Table 1 for sample sizes.

**Table 3:** Results of the LMMs on yolk THs and iodine in eggs from 1^st^ and 2^nd^ clutches laid by rock pigeon females treated with an I- or I+ diet.

| Predictor | Ln-Iodine |  |  | Ln-T<sub>4</sub> |  |  | T<sub>3</sub> |  |  |
| --- | --- | --- | --- | --- | --- | --- | --- | --- | --- |
|  | Estimate<br>(SE) | F<sub>ddf</sub> | p | Estimate<br>(SE) | F<sub>ddf</sub> | p | Estimate<br>(SE) | F<sub>ddf</sub> | p |
| Treatment (I+) | 2.04<br>(0.15) | 183.82<sub>21.5</sub> | **<0.001** | 0.05<br>(0.08) | 0.36<sub>21.3</sub> | 0.56 | 0.28<br>(0.18) | 2.40<sub>18.4</sub> | 0.14 |
| Exposure duration | 0.001<br>(0.002) | 0.14<sub>47.3</sub> | 0.71 | -0.001<br>(0.001) | 0.49<sub>44.6</sub> | 0.49 | 0.01<br>(0.005) | 7.59<sub>52.3</sub> | **0.01** |
| Complete clutch<br>(Yes) | -0.03<br>(0.13) | 0.07<sub>60.4</sub> | 0.80 | -0.04<br>(0.07) | 0.38<sub>57.7</sub> | 0.54 | -0.15<br>(0.20) | 0.54<sub>49.2</sub> | 0.47 |
| Treatment (I+) *<br>exposure duration | 0.007<br>(0.005) | 1.92<sub>46.1</sub> | 0.17 | 0.001<br>(0.003) | 0.11<sub>43.2</sub> | 0.74 | 0.01<br>(0.01) | 1.53<sub>49.2</sub> | 0.22 |
| Treatment (I+) *<br>complete clutch<br>(Yes) | 0.15<br>(0.26) | 0.33<sub>58.5</sub> | 0.57 | -0.05<br>(0.14) | 0.11<sub>56.4</sub> | 0.75 | 0.27<br>(0.40) | 0.45<sub>51.3</sub> | 0.51 |
| Clutch order (2) | -0.02<br>(0.07) | 0.11<sub>44.8</sub> | 0.74 | -0.02<br>(0.04) | 0.40<sub>42.4</sub> | 0.53 | 0.34<br>(0.13) | 6.84<sub>48.6</sub> | **0.01** |
| Treatment (I+) *<br>clutch order (2) | 0.007<br>(0.19) | 0.43<sub>43.8</sub> | 0.51 | 0.03<br>(0.07) | 0.11<sub>41.4</sub> | 0.74 | 0.42<br>(0.25) | 2.61<sub>47.0</sub> | 0.11 |

Yolk T_3_ was not affected by iodine treatment (EMMs (SE) T_3_: I- = 2.65 (0.19) pg/mg yolk, I+ = 2.89 (0.18) pg/mg yolk), but showed a slight increase over the exposure duration and with clutch order (Table 3; Fig. 5A). Yolk T_4_ was not affected by iodine treatment (back-transformed EMMs (SE) T_4_: I- = 7.10 (0.58) pg/mg yolk, I+ = 7.37 (0.55) pg/mg yolk), exposure duration of the treatment, or by clutch order (Table 3; Fig 5B). Finally, there were no effects of clutch completeness and its interaction with treatment on yolk THs (Table 3).

**Figure 5:**
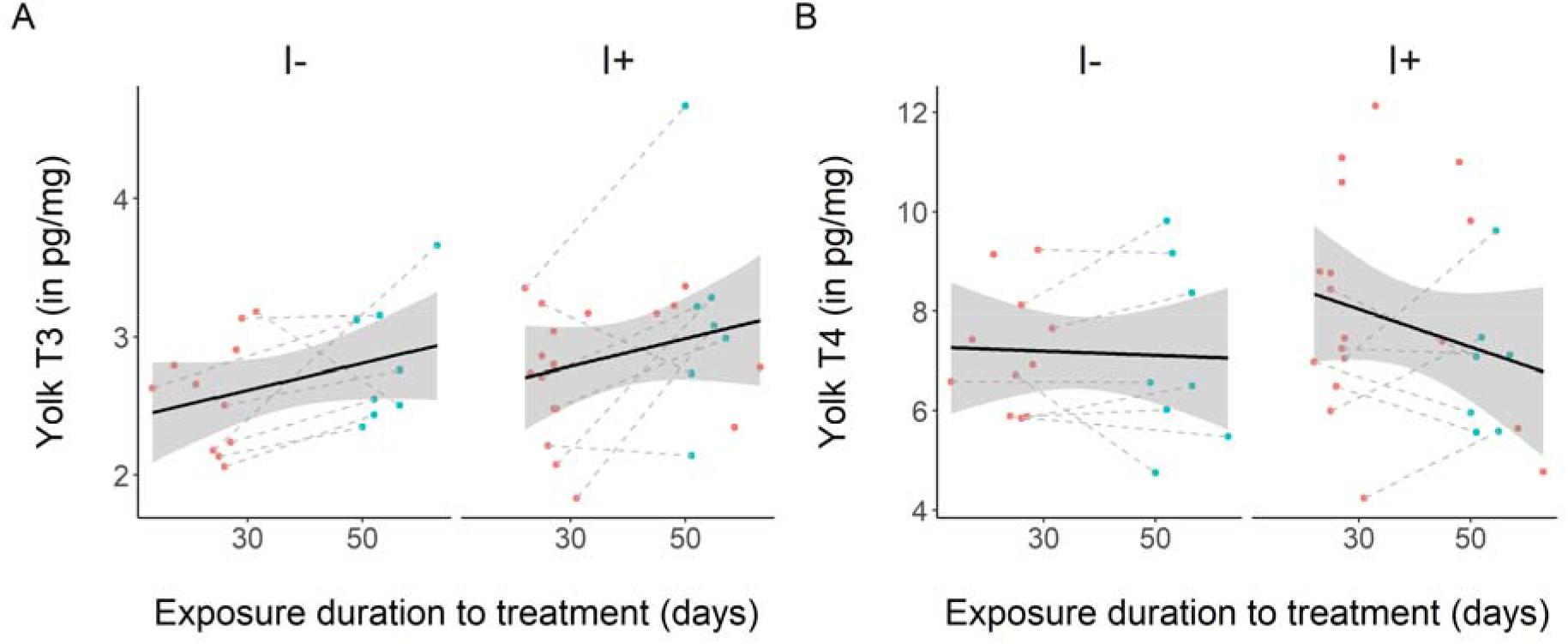
Yolk T_3_ (A) and T_4_ (B) in eggs from 1^st^ and 2^nd^ clutches laid by rock pigeon females treated with an I- or I+ diet. Red dots refer to eggs from the first clutches and blue dots refer to eggs from the second clutches. Black lines and shadow areas represent average and s.e.m values within each group, and grey dashed lines connect eggs from the same females. See Table 1 for sample sizes.

### Egg production

Overall, fewer females in the I- group laid their first clutches (complete or incomplete) than those in the I+ group (I- = 11 out of 19 females, I+ = 18 out of 19 females; χ^2^ = 5.24, p = 0.02; Table 1). This resulted in fewer eggs produced in the I- group than in the I+ group (I- = 17 eggs, I+ = 30 eggs; χ^2^ = 8.03, p = 0.005; Table 1). Focusing on the complete clutches, half as many females in the I- group laid complete clutches compared to those in the I+ group, yet difference was not statistically significant (I+ = 12/19 females, I- = 6/19 females; χ^2^ = 2.64, p = 0.10; Table 1).

## Discussion

In this study we tested whether dietary iodine limits mothers’ circulating TH concentration, TH transfer to the yolk, and egg production. To our knowledge, our study is the first to investigate the potential trade-off between circulating and yolk THs induced by low dietary iodine. We found that fewer females laid first clutches under the iodine-restricted diet compared to the females under the iodine-supplemented diet, resulting in a lower total number of eggs laid. We found that the iodine restricted diet decreased circulating and yolk iodine levels, though circulating and yolk THs were unaffected. Longer exposure to restricted iodine had no clear effect on circulating or yolk iodine and THs. Finally, we observed a slight increase in plasma T_4_ and in yolk T_3_ across time that was unrelated to the dietary iodine and is likely explained by seasonal changes or clutch order effects. Yet, because exposure duration to the treatment and clutch order are partly confounded, our experimental design does not allow us to fully disentangle both variables.

### Does restricted iodine induce a cost and a trade-off between circulating and yolk THs?

Our iodine-restricted diet successfully decreased circulating iodine concentrations compared with the supplemented diet. Despite this effect, we observed no differences in circulating TH concentrations. This is consistent with a previous study that showed that Japanese quails under limiting iodine availability maintained normal circulating THs concentrations (McNabb et al., 1985a). However, a similar study on ring doves found a decrease in circulating T_4_ concentrations with no changes in circulating T_3_, suggesting increased peripheral conversion from T_4_ to T_3_ to maintain normal T_3_ levels (McNichols and McNabb, 1987). The causes for discrepancies between our study and that of the dove study (McNichols and McNabb, 1987) are not clear. One potential explanation is that, in our study, we could only sample the females that laid eggs and thus apparently managed to maintain normal circulating THs despite restricted iodine whereas those that did no lay eggs may have suffered from low circulating TH concentrations.

In the yolk, like in the circulation, restricted dietary iodine decreased yolk iodine but not yolk TH concentration, in contrast to our predictions. The result of decreased yolk iodine is in line with a previous study on quails, which found that mothers fed with low dietary iodine produced eggs with low iodine but did not report yolk TH concentrations (McNabb et al., 1985a). Low egg iodine concentration in turn disturbs thyroid function in embryos and hatchlings (McNabb et al., 1985b; Stallard and McNabb, 1990). Circulating TH concentrations of embryos, however, were not affected by low egg iodine (McNabb et al., 1985b; Stallard and McNabb, 1990).

Contrary to previous studies that manipulated dietary iodine, we found that limited iodine availability hampered egg production, with 40% fewer females producing eggs in the I- group compared to the I+ group. However, females that managed to maintain normal circulating THs were also able to lay eggs with normal yolk THs, similar to the study by McNabb and colleagues (1985a). At the moment it is unclear why some females were affected and others not, but a potential explanation may be for example individual differences in the ability to store iodine. Two other studies found that administration of methimazole, a TH-production inhibitor, ceased egg laying in Japanese quails (Wilson and McNabb, 1997), and reduced egg production in chickens (Van Herck et al., 2013). These results suggest that our restricted diet might have induced hypothyroidism in some females, thus preventing them from laying eggs.

Therefore, we did not show evidence for a cost of restricted iodine in females that managed to lay eggs. Those females did not appear to face any trade-off between allocating iodine and THs to either self or their eggs. Yet, 40% of the females under the restricted diet paid a cost in terms of egg production. Whether this effect is due to limited production of THs is as yet unclear, but these females may have faced a trade-off between maintaining normal circulating THs or yolk TH deposition.

### Is there a regulatory mechanism to cope with the cost of restricted iodine and its associated trade-off?

The absence of an effect of restricted iodine on circulating and yolk THs suggest that mothers are not able to regulate yolk TH deposition independently from their own circulating THs, as recently proposed by Sarraude et al. (2020b). This is contradictory to previous studies showing evidence of independent regulation (Van Herck et al., 2013; Wilson and McNabb, 1997). However, the latter two studies induced supraphysiological hypo- or hyperthyroidism to the birds, which may explain such discrepancies.

Interestingly, we found that our restricted diet reduced egg production. As discussed above, this may be due to a hypothyroid condition that prevented females from laying eggs. This may have evolved to protect embryos from exposure to too low iodine and/or TH concentrations. In breeding hens, restricted dietary iodine can decrease egg hatchability (Rogler et al., 1959; Rogler et al., 1961b) and retard embryonic development (McNabb et al., 1985b; Rogler et al., 1959; Rogler et al., 1961b), and can induce thyroid gland hypertrophy in embryos and hatchlings (Rogler et al., 1961a; McNabb et al., 1985b; but see Stallard & McNabb, 1990).This may suggest that females may be able to regulate egg production when iodine availability is low. Overall, our results suggest that mothers in the restricted group appear to prioritise self-maintenance and offspring quality over offspring quantity.

### Restricted iodine and trade-offs in wild populations

Our low-iodine diet (0.06 mg I/kg food) is comparable to what birds may sometimes experience in the wild. Although relevant data are scarce, estimates of iodine content in food items such as barley and maize grains, wheat, or rye is highly variable, ranging from 0.06 to 0.4 mg I/ kg (Anke, 2004). Insectivorous species may also encounter iodine deficiency as the iodine content in insects vary from <0.10 up to 0.30 mg I/kg (Anke, 2004). As such low iodine availability can also be found in the wild, it is therefore relevant to study whether mothers may face trade-offs in iodine or TH allocation during the breeding season, or whether it influences egg laying itself. Yet, our study did not show evidence for the existence of a trade-off between circulating and yolk THs when environmental iodine is limited. Nevertheless, mothers may face a trade-off between allocating resources to themselves or producing eggs of sufficient quality.

In conclusion, we found that restricted dietary iodine did not decrease circulating or yolk THs despite reduced circulating and yolk iodine. Nevertheless, we found evidence that restricted availability of iodine induces a cost on egg production. Thus, mothers may not be able to regulate yolk TH transfer, but may be able to regulate egg production when facing limited iodine. Our results also indicate that females under limited iodine availability may prioritise their own metabolism over reproduction, or avoid exposing their offspring to detrimentally low iodine and/or THs. These explanations serve as interesting hypotheses for future research to further explore the consequences of limited iodine in wild populations.

## List of symbols and abbreviations

I: iodine
I-: iodine restricted diet
I+: iodine supplemented diet
T_3_: triiodothyronine
T_4_: thyroxine
THs: thyroid hormones

## Acknowledgments

We thank Martijn Salomons and the animal caretakers for helping us maintaining the colony. We are also grateful to Gerard Overkamp and Asmoro Lelono for their help with blood sampling. We also thank Bonnie de Vries and Tiphaine Bailly for their help with egg dissection.

## Funding

The study was funded by the Academy of Finland (grant no. 286278 to SR), the Ella and Georg Ehrnrooth Foundation (grant to BYH) and the University of Groningen (grant to TG). The funders had no role in study design, data collection and analysis, decision to publish, or preparation of the manuscript.

## Ethics

All the procedures were approved by the Centrale Commissie Dierproeven (AVD1050020185444) and the Animal Welfare Body of the University of Groningen (18544-01-001).

